# Characterisation of putative retrotrapezoid nucleus (RTN) chemoreceptor neurons in the adult human brainstem

**DOI:** 10.1101/2025.04.30.650987

**Authors:** Yazhou Liu, Rita Machaalani, Irit Markus, Claire E. Shepherd, Natasha N. Kumar

## Abstract

The retrotrapezoid nucleus (RTN) of rodents is located ventral to the facial motor nucleus (7N) and consists of acid-sensitive neurons that activate breathing and mediate the central component of the ventilatory response to hypercapnia. In rodents, RTN neurons can be histologically identified by the presence of paired-like homeobox 2B positive nuclei (Phox2b+) and the absence of cytoplasmic choline acetyltransferase (ChAT−) and tyrosine hydroxylase (TH−). Up to 50% of rodent RTN neurons synthesise galanin, and 88% express pituitary adenylate cyclase activating polypeptide (PACAP). The human RTN (hRTN) has not been mapped to date. This study aimed to map the location and cytoarchitecture of the adult hRTN and compare the findings to the homologies of rodents, macaques and human infants. Formalin-fixed, paraffin-embedded tissue blocks from two adult cases, spanning the medulla-pons, were serially sectioned (10µm thick) and every four in thirty sections was assayed for immunohistochemistry for ChAT, or double-labelled Phox2b/TH, Phox2b/galanin and Phox2b/PACAP, followed by analysis using QuPath software. hRTN neurons, identified as Phox2b+/TH−/ChAT−, were located ventral to 7N and lateral to the superior olive, overlapped with the C1 or A5 catecholaminergic population and extended rostrocaudally from Obex +13 to +17 mm. In the parafacial area, 90% of Phox2b immunoreactive (−ir) neurons are hRTN neurons, totaling around 5000 bilaterally, and were surrounded by numerous TH-ir fibers. Galanin- and PACAP-ir was identified in 43% and 39% of Phox2b-ir parafacial neurons, respectively. This is the first study to characterise and quantitatively map the adult human RTN using a series of neurochemical markers.

## Introduction

In mammals, breathing is regulated by central respiratory nuclei and a neural network in the brainstem (Ikeda et al., 2017). Central respiratory chemoreceptors are clusters of cells in the lower brainstem respiratory network which regulate breathing stimulated by carbon dioxide (CO_2_) or its proxy, protons (H^+^). Acid sensing chemoreceptor neurons in the retrotrapezoid nucleus (RTN) (Guyenet and Bayliss, 2015, Guyenet and Bayliss, 2022) project to the ventral respiratory column (VRC) which drives the central respiratory chemoreflex response (Guyenet et al., 2019). Impaired central respiratory chemoreception results in an impaired hypercapnic ventilatory response, as occurs in congenital hypoventilation syndrome (CCHS) and central sleep apnoea (Kasi et al., 2018, Meylemans et al., 2021, Degl’Innocenti et al., 2018, Amiel et al., 2009).

The RTN comprises about 2000 neurons in the rat, and 700 - 800 neurons in mice, bilaterally (Takakura et al., 2008, Guyenet and Bayliss, 2015, Shi et al., 2017, Lazarenko et al., 2009). Phox2b is the most widely recognised biomarker for RTN chemoreceptors, and changes in its expression are regarded as an indicator of RTN chemoreceptor dysfunction associated with various central respiratory disorders in both rodents and humans (Amiel et al., 2009, Goridis and Brunet, 2010, Meylemans et al., 2021, Lavezzi et al., 2012). In addition to Phox2b, alterations in neuropeptides, such as galanin and PACAP, are important for regulation of central chemoreception in the RTN. In rodents, galanin is expressed in approximately 50% of RTN neurons (Stornetta et al., 2009) and the PreBotzinger complex receives glutamatergic innervation from galaninergic RTN neurons (Bochorishvili et al., 2012, Dereli et al., 2024). Long term hypercapnia induces an increase in preprogalanin (PPGAL) expression in the RTN (Dereli et al., 2019) and microinjection of galanin into the VRC, specifically in the PreBotzinger and Botzinger complexes, impairs the chemoreflex response to hypercapnia (Abbott et al., 2009). In neonatal mice, PACAP is expressed in 100% of RTN chemoreceptor neurons and the deletion of PACAP in RTN neurons leads to elevated incidence of apnoea’s and reduced respiratory responses to CO_2_ stimulation in the postnatal period (Shi et al., 2021). Although PACAP expression in the human brainstem has been studied (Huang et al., 2017), the RTN was not investigated due to the lack of knowledge regarding its precise location.

To date, four studies have investigated the location of the RTN in primates; 1 in monkeys and 3 in humans (Table 1). Levy and colleagues aimed to identify Phox2b mRNA and protein in formalin-fixed, paraffin-embedded (FFPE) adult human brainstem, but it failed to be detected and as such, the data was on a population of PPGAL/ Solute Carrier Family 17 Member A6 (SLC17A6) mRNA containing neurons in the parafacial region (Levy et al., 2021). The other two studies were conducted on FFPE tissue from foetuses or infants (Rudzinski and Kapur, 2010, Lavezzi et al., 2012). Rudzinski and Kapur (2010), described the putative human RTN (hRTN) as a discontinuous cluster of small- to medium-sized Phox2b positive/ Neurokinin 1 receptor positive/Tyrosine hydroxylase negative (Phox2b+/NK1R+/TH−) neurons situated ventral to the facial motor nucleus (7N) and lateral to the superior olivary nucleus (SO) spanning from the caudal pons to the rostral medulla (Rudzinski and Kapur, 2010). Lavezzi et al (2012) described the RTN as a cluster of Phox2b immunoreactive (−ir) neurons situated between the 7N and SO in the caudal pons (Lavezzi et al., 2012). While both infant studies sought to characterise the hRTN, they were restricted by limited tissue sampling and inconsistent Phox2b-ir patterns (Table 1).

**Table 1:**
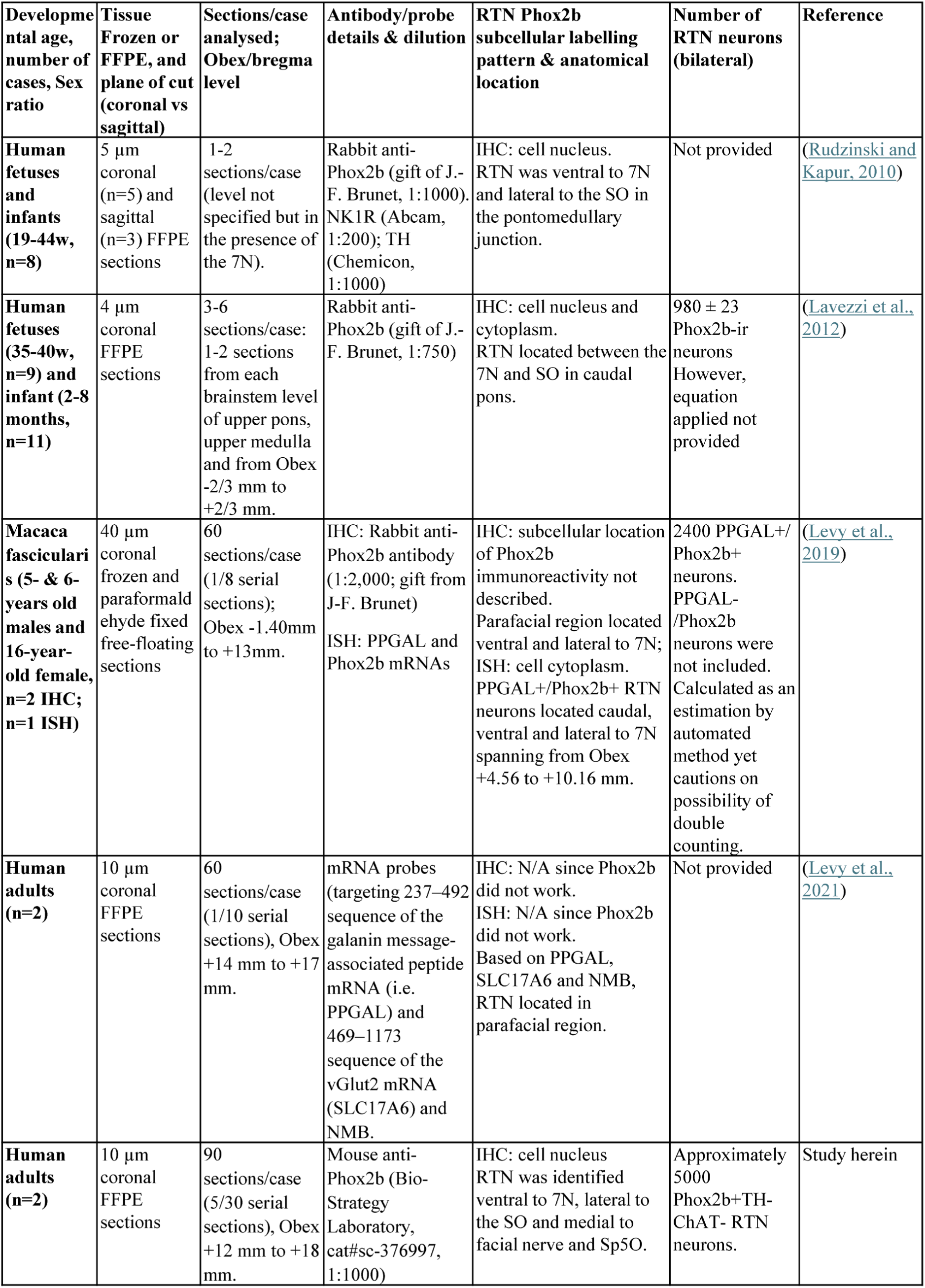
Summary of data collected on primate RTN in chronological order.

Since the RTN has not been systematically mapped using Phox2b in the human brainstem, our overall study goal was to fill this research gap. This immunohistochemical study used post-mortem FFPE tissue from 2 adult humans to: 1) characterize Phox2b-ir in the RTN region, 2) establish the neurochemical signature of hRTN neurons and quantify these neurons, 3) establish the neuroanatomical location and dimensions of the hRTN relative to 7N, C1 or A5 catecholaminergic neurons, 4) determine the proportion of parafacial Phox2b-ir neurons expressing the neuropeptides galanin and PACAP, and 5) describe the morphological features (size and density) of parafacial Phox2b-ir neurons. Our findings provide valuable insights for investigations into Phox2b-associated disorders, particularly those involving respiratory and autonomic dysfunction including CCHS and sleep apnoea.

## Methods

### Human Tissue Selection

Post-mortem, formalin-fixed brainstem specimens from two adult human cases (76-year-old female, and 87-year-old male; Supplementary Table 1), were sourced from the Sydney Brain Bank through regional brain donor programs in Sydney, Australia. Specimens were acquired from asymptomatic individuals exhibiting solely age-related brain changes and devoid of significant neuropathology.

Exclusion criteria included documented respiratory drive disorders, post-mortem delays exceeding 72 hours and formalin fixation exceeding ten years. The demographic and autopsy characteristics are presented in Supplementary Table 1.

Brainstem tissue blocks spanning from the rostral medulla oblongata (Obex +12 mm) to the caudal pons (Obex +18 mm) were selected (Fig. 1), as this region contains the 7N and is expected to include the hRTN. This Obex range was guided by a human brainstem atlas (Paxinos et al., 2020).

**Fig. 1.**
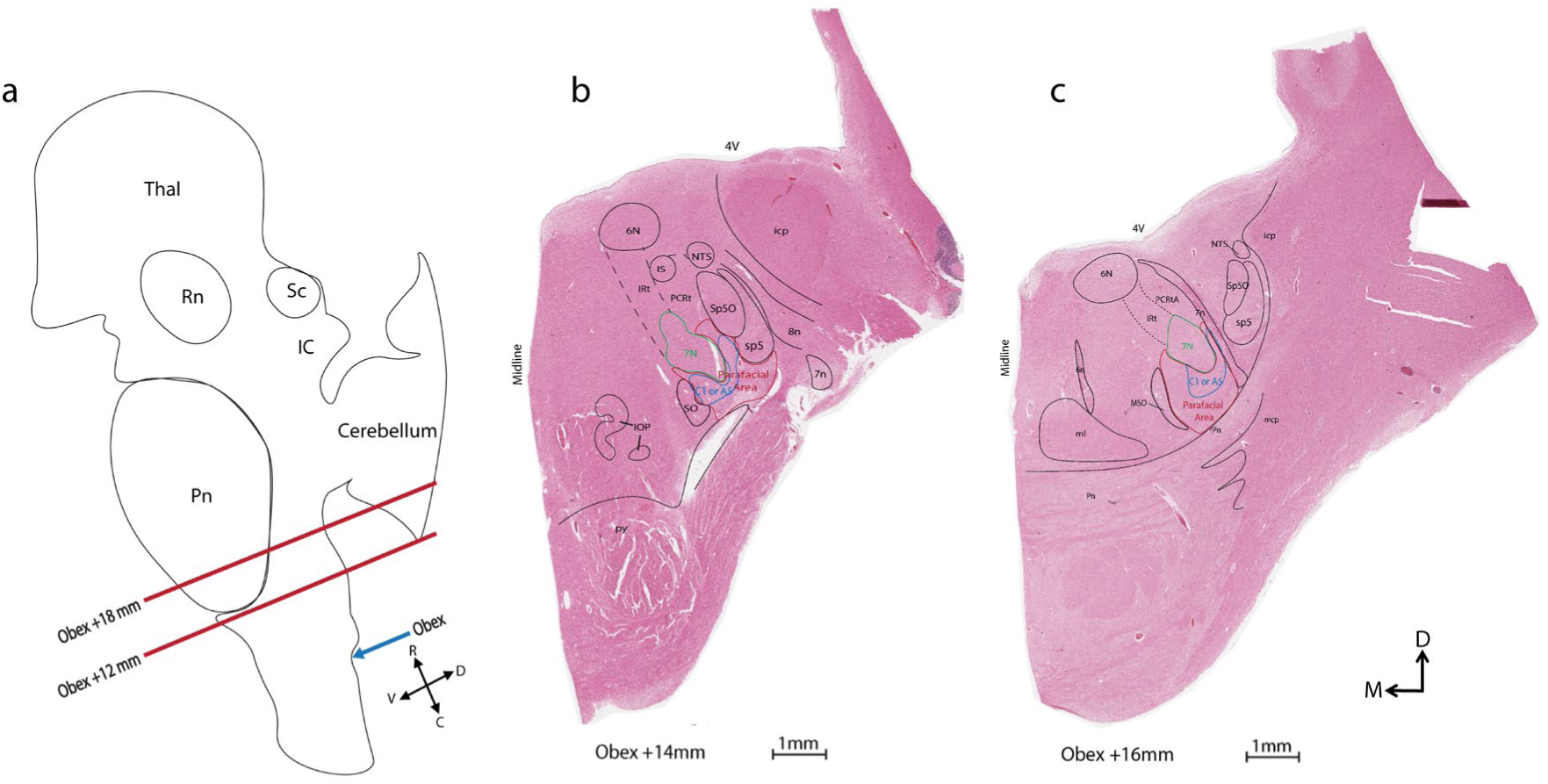
Tissue sectioning and parafacial area localisation. (a) Sagittal schematic of the adult human brainstem and thalamus indicating the angle of tissue sectioning and location spanning from Obex (indicated by blue arrow) +12 to +18 mm (between red lines). (b & c) Representative H&E stained caudal (Obex +14 mm) and rostral (Obex +16 mm) hemi-sections overlaid with brainstem schematics. (b) Caudally, the parafacial (red outline) is encapsulated by a region medial to sp5 and Sp5O, ventral to 7N, lateral to the SO and overlapping with C1 or A5 neurons (blue outline). (c) Rostrally, the parafacial area is medial to 7n, ventral to 7N, lateral to MSO and overlaps with C1 or A5 neurons (blue outline). Adapted from (Paxinos et al., 2020)

### Tissue sectioning

Ten-micron serial sections were cut from 2-3 mm thick transverse oriented FFPE brainstem blocks, using a rotary microtome. We collected approximately 700 sections per case, extending rostrocaudally from Obex +18 to +12 mm (Fig. 1). One in every thirty serial sections (300 μm apart) was stained with Hematoxylin & Eosin (H&E), to obtain representative sections containing the 7N. After mapping the full extent of the 7N in H&E sections, four of every thirty serial sections were selected for immunohistochemistry using antibodies against ChAT, Phox2b+TH, Phox2b+PACAP and Phox2b+galanin, respectively (Supplementary Table 2).

### Immunohistochemistry (IHC) Protocol for Human FFPE Brain Tissue

The IHC protocol was adapted from our previous methods (Huang et al., 2017), with necessary optimisations for Phox2b detailed in Supplementary Material 1. Antigen retrieval was performed using heat-induced epitope retrieval (HIER) with citrate buffer (pH 6, #S2369, Agilent DAKO) for 15 minutes. After a 2-hour block in 10% normal donkey serum (NDS) in tris buffered saline (TBS) with 0.3% Tween 20, sections were incubated overnight with goat anti-ChAT followed by a 2-hour incubation with biotinylated secondary antibody, in 2% NDS /TBS with 0.1% Tween 20 (Table 2). Signal amplification used the Vectastain Elite ABC HRP Kit (#PK-6100, Vector Laboratories), and visualisation was achieved using 3,3’-diaminobenzidine (DAB Substrate Kit, #SK-4100, Vector Laboratories).

**Table 2.**
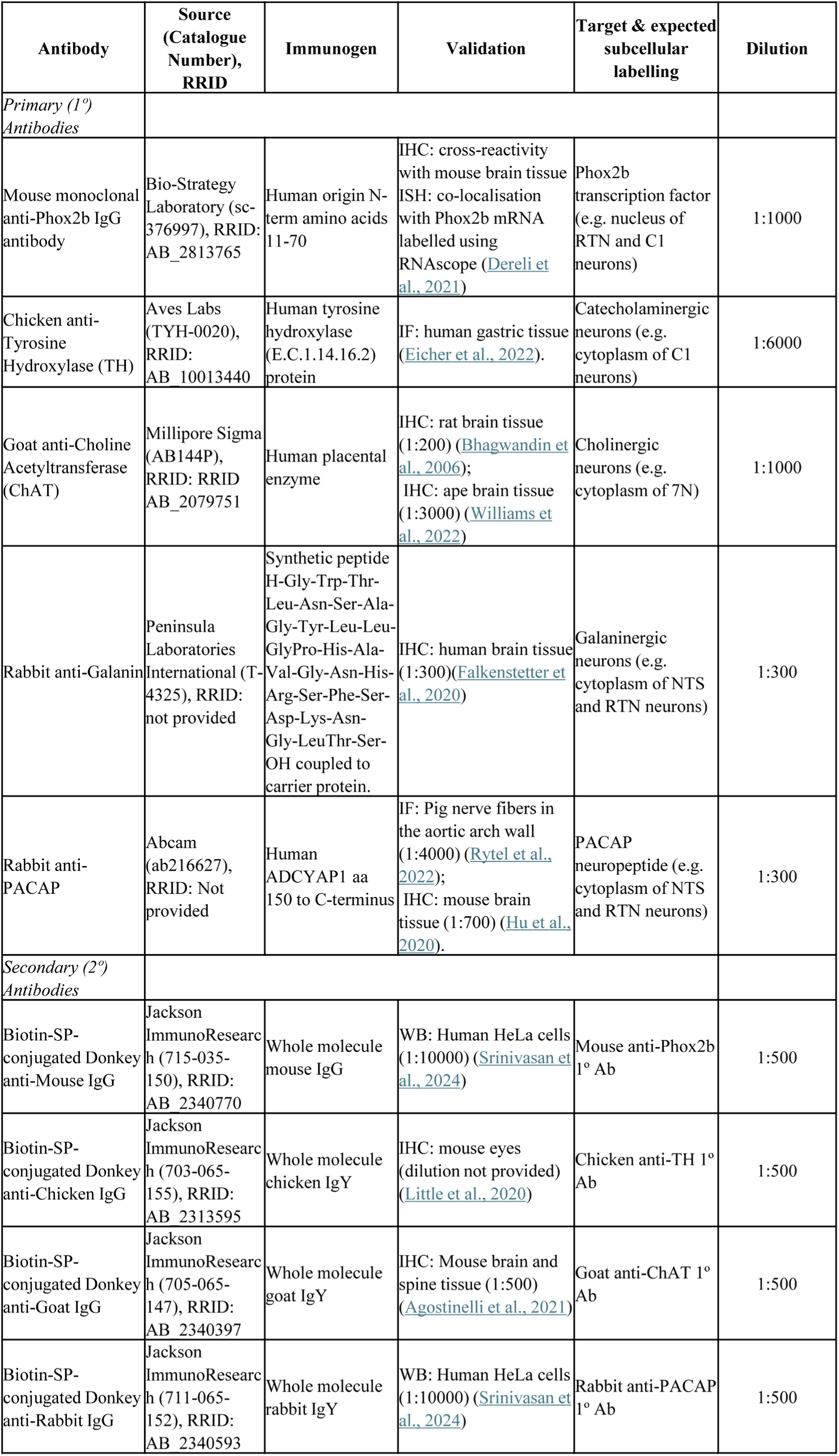
Primary and secondary antibodies used for immunohistochemistry (IHC)

For dual labelling (Phox2b+TH, Phox2b+PACAP, or Phox2b+galanin), Phox2b was detected first and visualised using DAB with nickel enhancement (black) (#SK-4100, Vector Laboratories). In the sequential round of immunohistochemistry, the tissue was incubated with a primary antibody against TH, PACAP, or galanin, followed by the corresponding secondary antibody and DAB (brown) visualisation. After dehydration and clearing, sections were coverslipped using dibutylphthalate polystyrene xylene (DPX). Antibody details and concentrations are listed in Table 2.

### Microscopy and imaging

Slides were digitised using a slide scanner (Aperio AT2, Leica BioSystems) at 20× magnification. The scanned files were viewed using QuPath Software (Version 0.2.3) (Bankhead et al., 2017). Coronal hemi-sections approximating from Obex +12 to +18 mm were assessed.

The parafacial area was defined as follows: rostrocaudally, spans from the rostral ventrolateral medulla to the caudal ventrolateral pons, located adjacent to the facial motor nucleus (Obex +13 to +17 mm),dorsoventrally and mediolaterally, it is ventral to the spinal trigeminal nucleus, oral part (Sp5O), dorsal to ventral medullary surface, lateral to the intermediate reticular nucleus (IRt) and inferior olive (IO), and medial to the facial nerve (7n) (Fig. 1). Schematic diagrams were drawn using Adobe Illustrator 2024 (Adobe Inc., San Jose, CA, USA).

### Quantitative analyses

The cell diameter, and area of the cell soma and nucleus of RTN neurons, were measured using QuPath tools (See details in Supplementary Material 2). This data was collated using Microsoft Excel v14.0 (2010, Microsoft, USA), and exported into GraphPad Prism (Version 9, CA, USA).

Neurons with the following neurochemical signatures were counted:

1. Phox2b immunoreactivity (−ir) parafacial neurons, located ventrolateral to the 7N, as:

a. Phox2b-ir RTN neurons: Phox2b+ nucleus, TH− and ChAT− cytoplasm,
b. PACAP+/Phox2b+ parafacial neurons and PACAP−/Phox2b+ parafacial neurons,
c. Galanin+/Phox2b− parafacial neurons and galanin−/Phox2b+ parafacial neurons,
2. 7N neurons as ChAT+ cytoplasm
3. C1 neurons: Phox2b+ nucleus, TH+ cytoplasm (Kang et al., 2007, Agostinelli et al., 2023),
4. A5 neurons: Phox2b− nucleus, TH+ cytoplasm (Kang et al., 2007, Agostinelli et al., 2023).

Phox2b-negative RTN neurons were not investigated in this study. The RTN neurons referred in this study were Phox2b-ir RTN neuron. Number of RTN neurons was calculated by applying the formula derived from (Abercrombie, 1946):

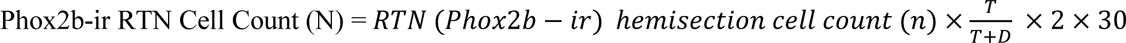

N, Total Phox2b-ir RTN cell count; n, RTN cell count/hemi-section; T, section thickness; D, average diameter of the cell.

Morphological data from the 2 adult cases was averaged and presented as mean ± standard deviation. No statistical analysis was performed due to the study being an anatomical description of the hRTN, without comparison between a control and experimental group.

## Results

### Immunoreactive labelling profile of antibodies used in this study

The subcellular localisation of labelling for each primary antibody was as expected in positive control regions, including the 7N, NTS and C1 (Fig. 2) (Huang et al., 2017, Kordower et al., 1992, Levy et al., 2019, Agostinelli et al., 2023). ChAT, TH, PACAP and galanin-ir were observed in the cytoplasm and fibers, while Phox2b-ir was confined to the nucleus of neurons (Fig. 2).

**Fig. 2.**
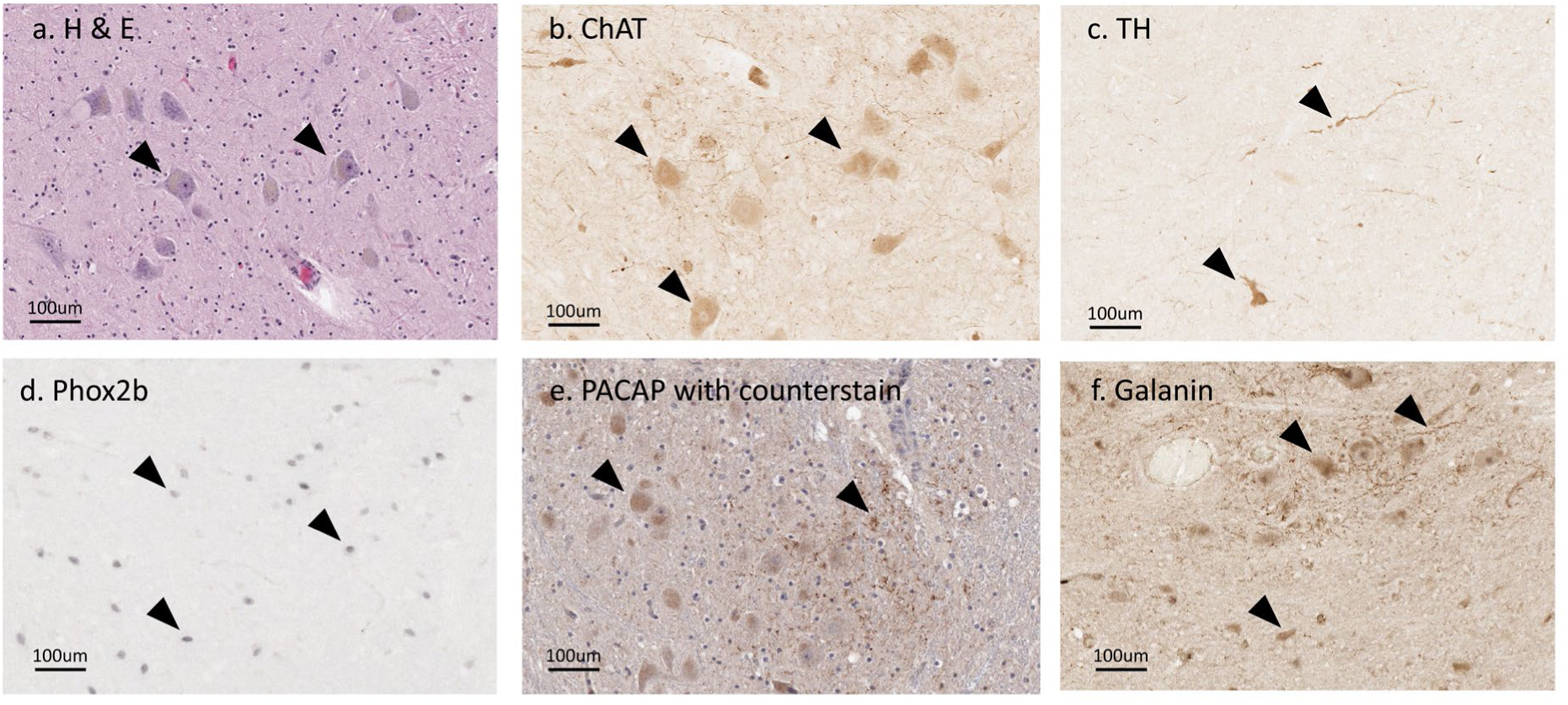
Exemplary H&E and primary antibody labelling. (a) H&E staining was performed to identify 7N neurons, based on their anatomical location. (b) ChAT-ir cytoplasm and fibers in motor neurons within the 7N. (c) TH-ir cytoplasm and fibers in catecholaminergic C1 neurons. (d) Phox2b-ir nuclei in NTS neurons. (e) PACAP-ir cytoplasm and fibers of hematoxylin counterstained NTS neurons. (f) Galanin-ir cytoplasm and fibers in the NTS. Arrowheads point to examples of positive immunoreactivity; arrowheads in (a) and (b) point to the same 7N cells

Phox2b-ir was subcategorised as either strong or weak (Supplementary Fig. 1), with cells in each category counted separately before totals were combined (see Supplementary Table 3).

### Neurochemical signature of putative hRTN neurons

Identification of RTN neurons according to their biochemical signature of Phox2b+ nucleus, TH−/ChAT− cytoplasm, was clearly discernable from neighboring 7N (ChAT+) and overlapping C1 or A5 (TH+) neurons (Fig. 3a, e, f and Supplementary Fig. 2). TH+ fibers coursed through the RTN region (Fig. 3a). In a high magnification image (Fig. 3b), the TH-positive axons were clearly delineated, and TH+ boutons made close appositions with RTN neurons. The subcellular labelling of these neurochemical markers in the RTN and nearby nuclei (7N, C1 or A5) is documented in Table 3. A few Phox2b+/TH− neurons were also detected in Sp5O (Fig. 3c), IRt and parvocellular reticular nucleus (PCRt) (Fig. 3d).

**Fig. 3.**
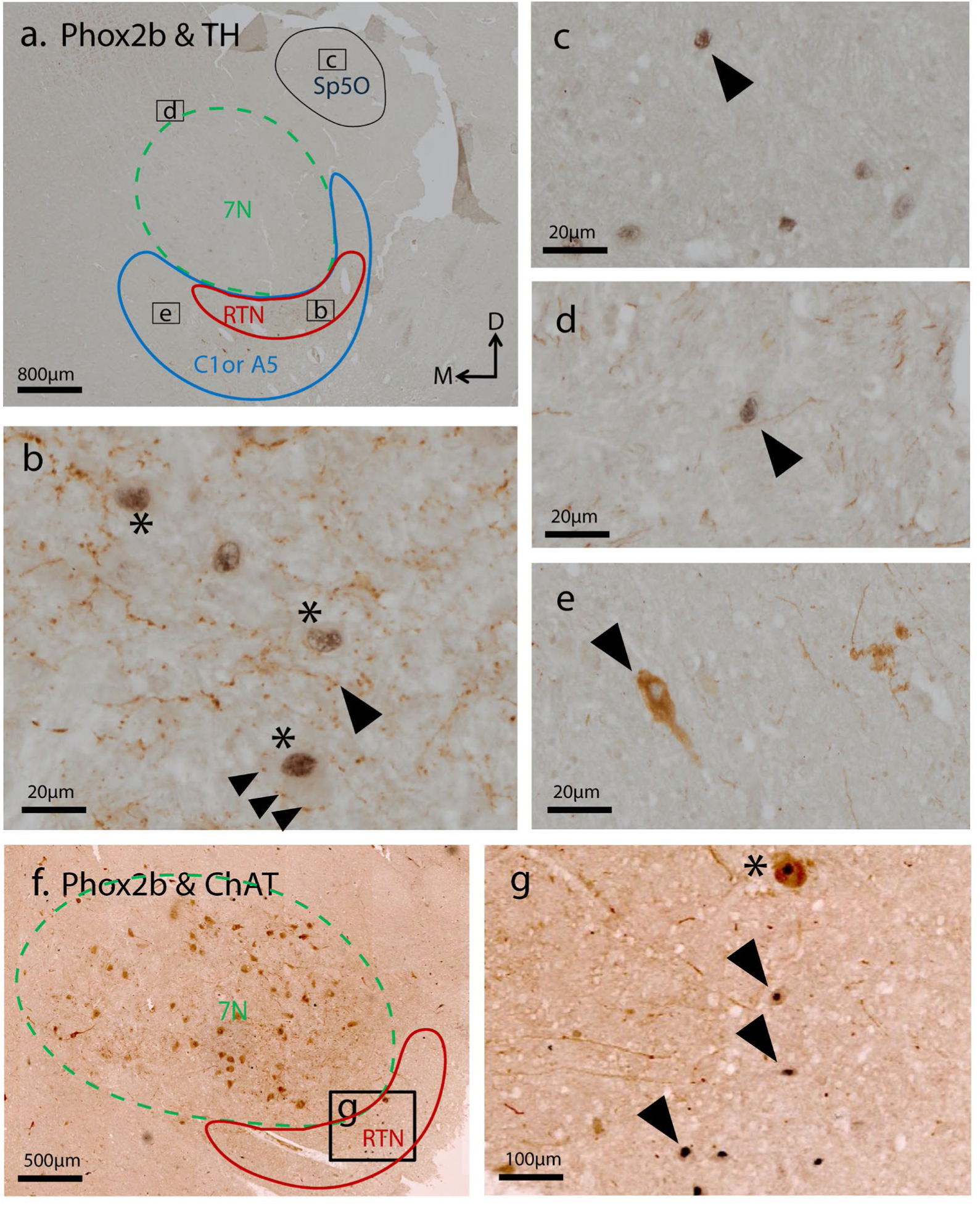
RTN neurochemical signature and spatial relationship to 7N, C1 or A5. (a) Low-magnification image showing the anatomical locations of the RTN (red outline), C1 or A5 region (blue outline), Sp5O (black outline), and 7N (green outline) within the brainstem. (b) Dual immunohistochemistry for Phox2b and TH identified RTN neurons as Phox2b+ with nuclear localisation (asterisks), and TH− cytoplasm, distinguishing RTN neurons from C1 or A5 neurons. Numerous TH-ir fibers (arrowheads) were present in the RTN region and closely apposed Phox2b-ir neurons (asterisks). (c) Sp5O neurons exhibited Phox2b-ir (arrowhead). (d) A few sparsely distributed Phox2b+/TH− neurons were detected in the IRt, dorsomedial to the 7N (arrowhead). (e) A5 neuron expressing TH-ir in the cytoplasm. (f) Low power image illustrating the location of the RTN (red outline) ventral to the 7N (green outline) in the brainstem (Obex +15 mm). (g) Phox2b and ChAT double immunohistochemical detection demonstrate RTN neurons as Phox2b+ and ChAT− (arrowheads). In contrast, the 7N neurons (asterisk) were identified by their ChAT+ soma and relatively large diameter

**Table 3.** Morphological characteristics of RTN, 7N, C1 and A5 neurons, based on their respective subcellular neurochemical signature.

| Cell Type | Morphology | Nucleus: Area ($\mu\text{m}^2$) | Cell Area ($\mu\text{m}^2$) | Cell Diameter ($\mu\text{m}$) | Subcellular neurochemical profile | | |
| --- | --- | --- | --- | --- | --- | --- | --- |
|  |  |  |  |  | Cytoplasm | Nucleus | Axon |
| 7N neurons (n=68, 1 section/case) | Large multipolar neurons with short dendrites | 274.6 ± 108 | 1194.0 ± 371.8 | 41.8 ± 10.7 | ChAT+, TH-, PACAP+/-, galanin- | Phox2b+/- | ChAT+ |
| RTN neurons (n=241, 3 sections/case) | Medium round or fusiform neurons with short dendrites | 69.0 ± 23.3 | 309.9 ± 61.8 | 19.9 ± 4.0 | ChAT-, TH-, PACAP+/-, Galanin+/- | Phox2b+ | Not detected |
| C1 neurons (n=38, 4 sections/case) | Medium multipolar neuron with short dendrites and long projecting axons | 98.0 ± 51.2 | 528.2 ± 165.2 | 35.1 ± 8.9 | ChAT-, TH+, PACAP+/-, galanin- | Phox2b+ | TH+ |
| A5 neurons (n=12, 4 sections/case) | Medium multipolar neuron with short dendrites and long projecting axons | 101.3 ± 44.9 | 468.7 ± 169.4 | 33.9 ± 11.7 | ChAT-, TH+, PACAP+/-, galanin- | Phox2b- | TH+ |

### Phox2b+ parafacial neurons are immunoreactive for galanin and PACAP

Although TH immunohistochemistry was not performed in combination with Phox2b/PACAP or Phox2b/galanin labelling, our quantitative data show that 90% of Phox2b-ir neurons located ventral to 7N are devoid of TH-ir (Fig. 3b). This strongly suggests that these neurons are part of the RTN population. Therefore, in this context, we confidently identified RTN neurons based on their Phox2b expression and anatomical location within the parafacial region. Based on the double labelling for Phox2b and galanin as well as the double labelling for Phox2b and PACAP, a total of 5295 Phox2b+ parafacial neurons were counted (Table 4).

**Table 4.**
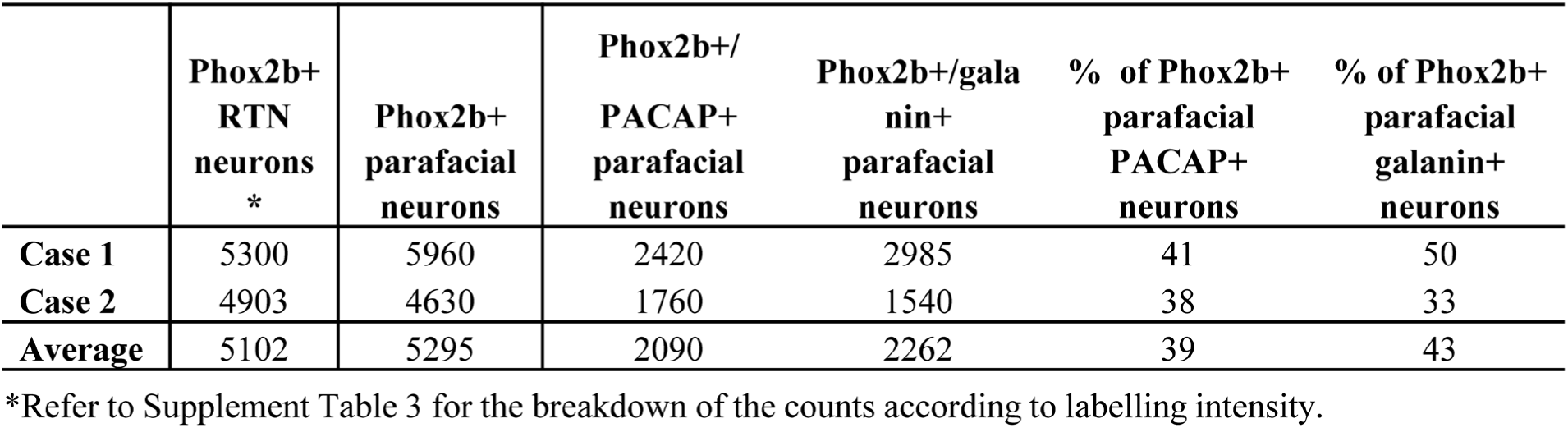
Total Phox2b-ir parafacial neurons and the proportion that are galanin-ir or PACAP-ir*.

Galanin-ir was detected in the soma and dendrites of Phox2b-ir parafacial neurons (Fig. 4a-c) and represented 43% of total Phox2b-ir population (2262 galanin+/Phox2b+ neurons out of 5295 Phox2b+ parafacial neurons) (n=2, with 14 sections counted per case) (Supplementary Table 3). Similarly, PACAP-ir was confined to cell soma and dendrites of 39% Phox2b-ir parafacial neurons (2090 PACAP+/Phox2b+ parafacial neurons out of 5295 Phox2b-ir parafacial neurons, Fig. 4d-f, Supplementary Table 3).

**Fig. 4.**
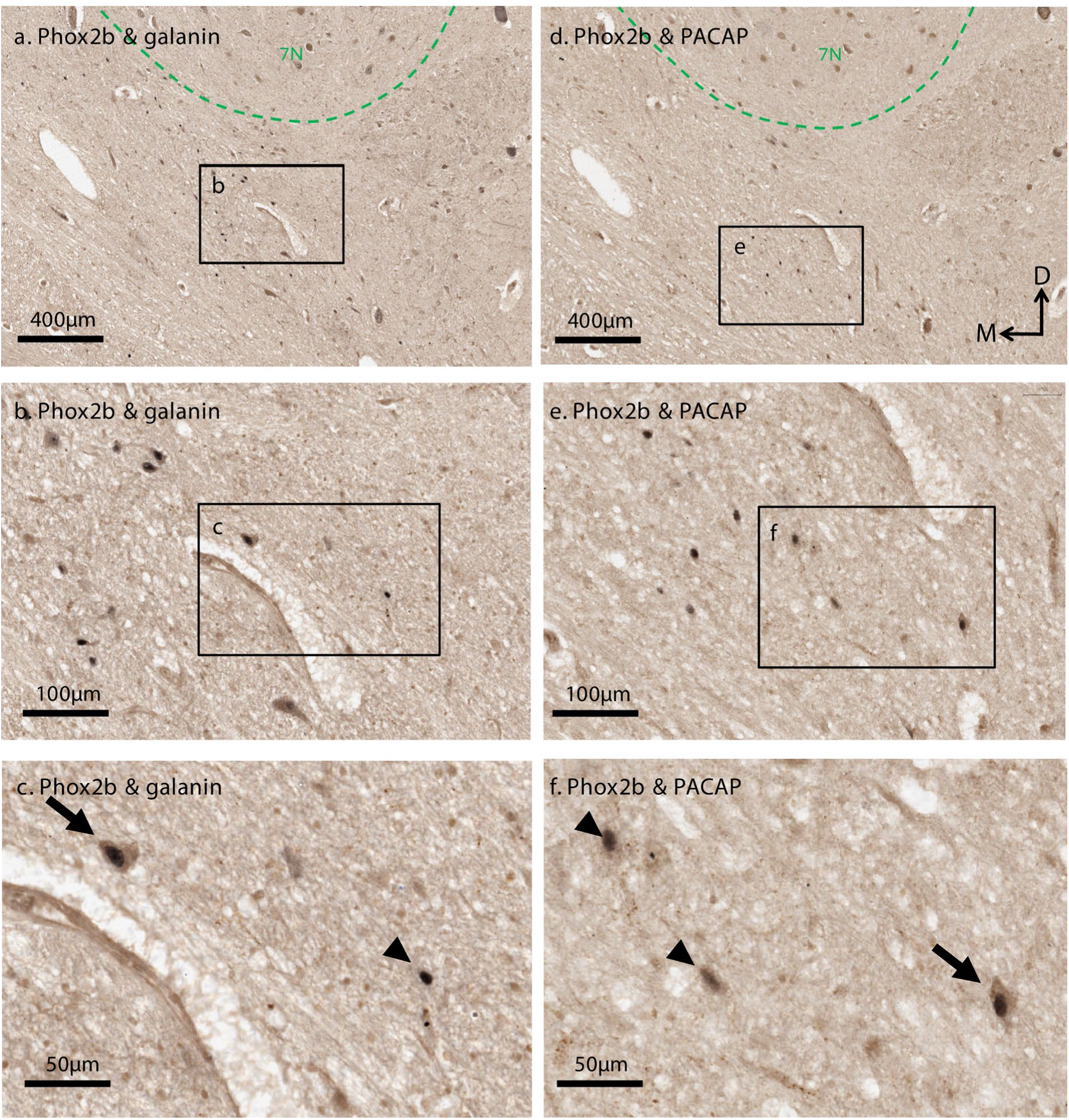
Galanin and PACAP immunoreactivities are detectable in large subsets of Phox2b-ir parafacial neurons. (a and d) Low magnification image of the RTN region immediately ventral to the 7N. (b) Many neurons in the parafacial region are both galaninergic (brown cytoplasmic labelling) and Phox2b-ir (black nucleus labelling). (c) Magnification of inset in b) demonstrating a galanin+/Phox2b+ neuron (arrow) and a galanin−/Phox2b+ neuron (arrowhead). (e) A subgroup of neurons in the parafacial region display PACAP-ir (brown cytoplasmic labelling) and Phox2b-ir (black nucleus labelling). (f) Magnification of inset in (e) demonstrating PACAP+/Phox2b+ neuron (arrow) and PACAP−/Phox2b+ neurons (arrowheads)

### Anatomical characteristics of the hRTN based on the distribution of Phox2b+/TH− neurons

Putative hRTN neurons were located ventral and ventrolateral to the 7N, lateral to the SO, dorsomedial to the pontomedullary junction, medial to the Sp5O, spinal trigeminal tract (sp5) and the facial nerve, and overlapping with C1 or A5 catecholaminergic neurons (Fig. 3).

RTN neurons were found from Obex +13 to +17 mm (Fig. 5 and Fig. 6) of the medulla oblongata. At caudal levels, Obex +13 to +13.8 mm, RTN neurons were sparsely distributed (Fig. 5 and Fig. 6a). By contrast, within the range of +13.8 to +14.8 mm, RTN neurons formed a distinct and prominent cluster (Fig. 5), indicated by a concentrated presence of Phox2b+/TH− neurons. More rostrally, (+14.8 to +16.8 mm), the RTN neurons were again more sparsely distributed, devoid of a distinct cluster (Fig. 5 and Fig. 6c). Anatomically, rostral RTN neurons were distributed medial to the facial nerve, ventrolateral to the 7N, dorsal to the ventral surface of medulla and lateral to the SO (Fig. 5).

**Fig. 5.**
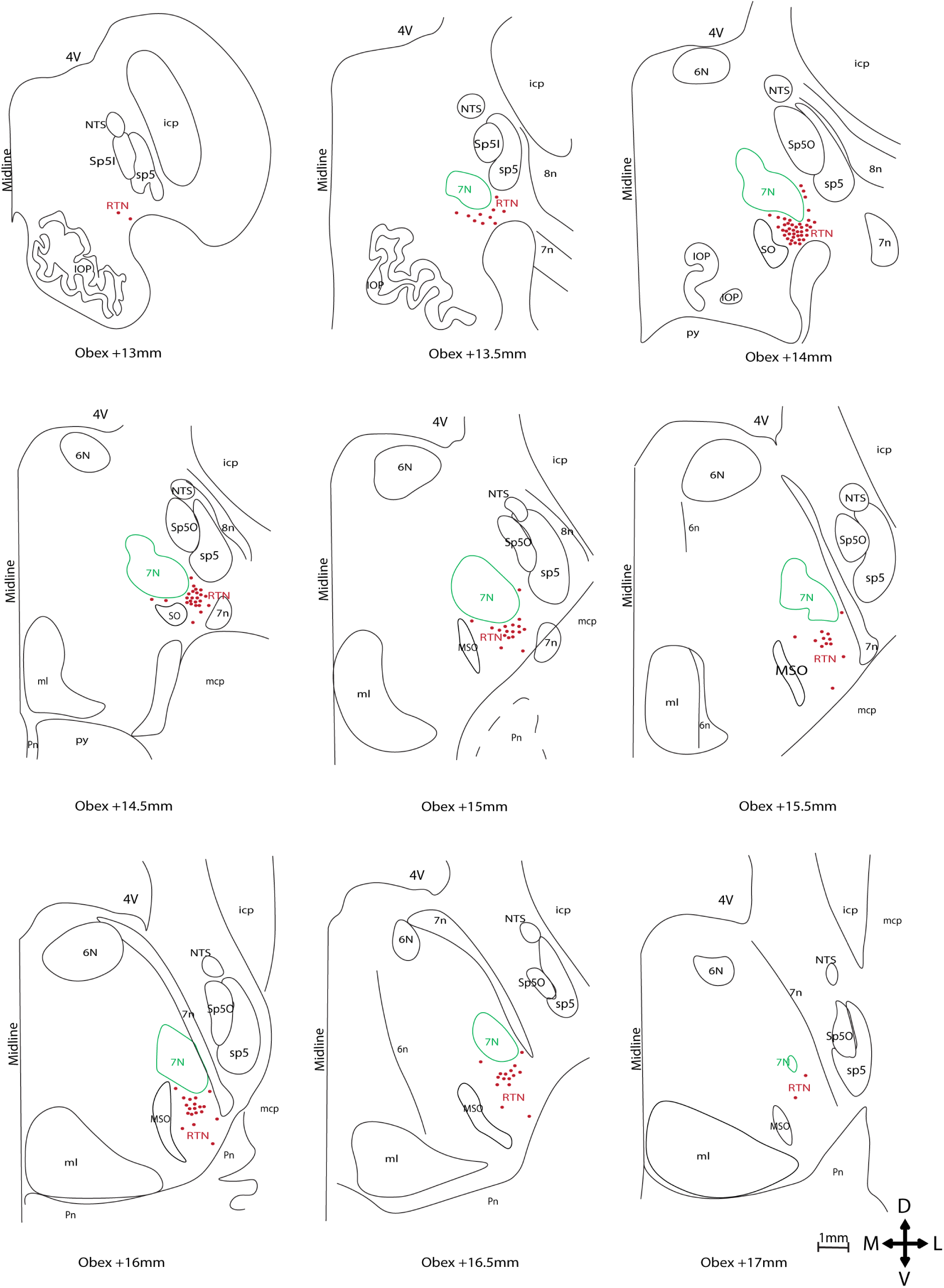
Schematic coronal hemi-sections illustrating the distribution of parafacial Phox2b+/TH− neurons, representative of the RTN (from Obex +13 to +17 mm). Red dots represent Phox2b+/TH− neurons. RTN neurons are located immediately ventral to the 7N, lateral to the SO, and medial to the facial nerve (rostrally) or spinal trigeminal nucleus (caudally). The most densely packed cluster of RTN neurons is situated 14 mm rostral to Obex. Drawings were made with reference to (Paxinos et al., 2020)

**Fig. 6.**
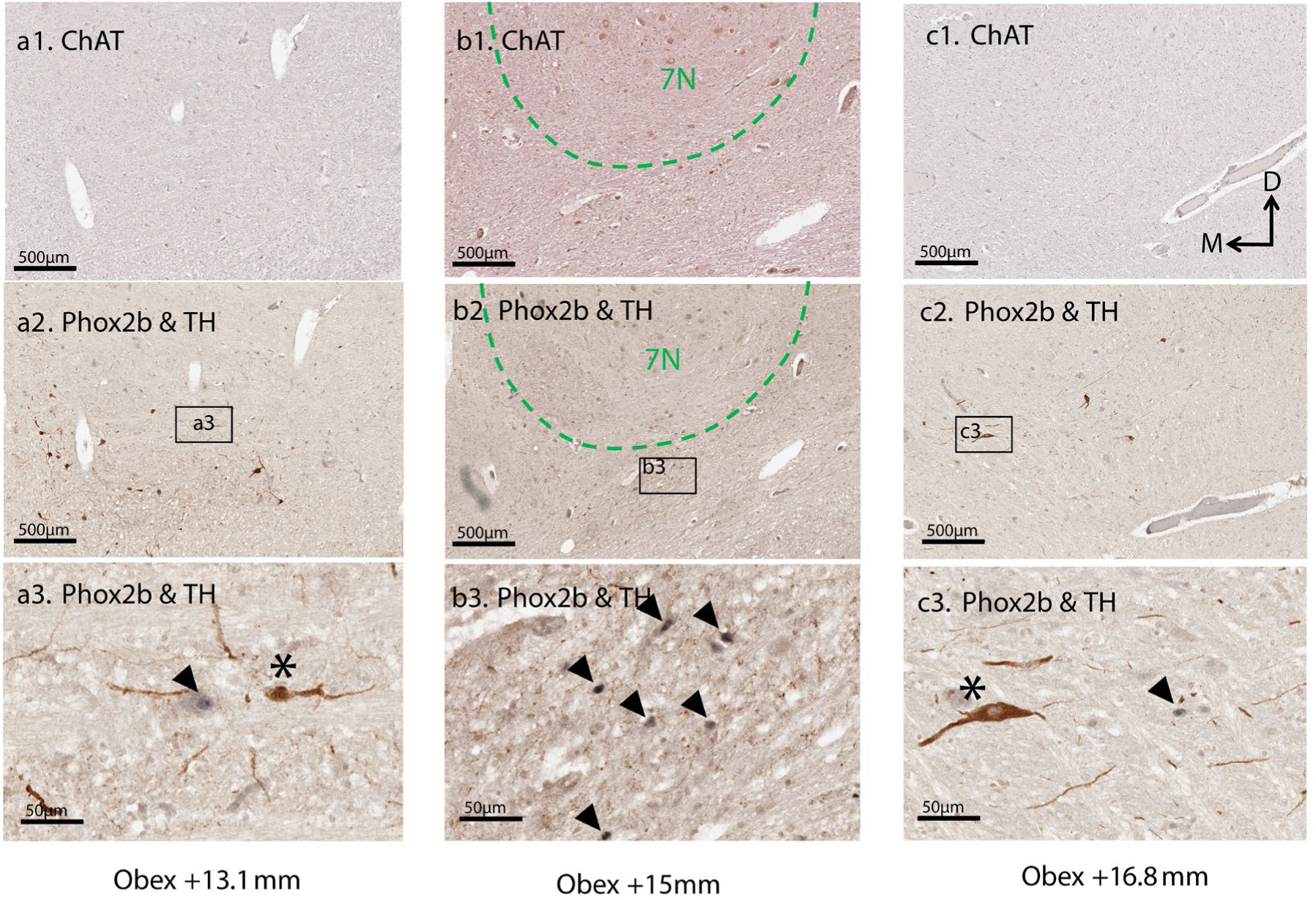
RTN neurons extend along the entire rostrocaudal length of the 7N. (a1 and a2) Caudal pole of the 7N (Obex +13.1 mm). (a3) C1 neuron (TH-ir, asterisk) next to an RTN neuron (arrowhead). (b1 and b2) 7N neurons are distinguished from the RTN region by the green outline. (b3) RTN neurons (Phox2b+/TH−, arrowheads) and TH-ir fibers (brown puncta) are notably abundant at this level of the RTN. (c1 and c2) Rostral pole of the 7N (Obex +16.8 mm). (c3) Few RTN neurons (Phox2b+/TH−, arrowhead) were present. An A5 neuron (Phox2b−/TH+, asterisk) is visible in the rostral RTN region

### Morphology of hRTN neurons

The soma of RTN neurons was typically round, or fusiform, with one or two dendrites extending in various directions (Fig. 4). RTN neurons were significantly smaller in size compared to 7N, C1 or A5 neurons, with an average soma area of 309.9 ± 61.8 µm² (n=241 cells), and an average nucleus area of 69.0 ± 23.3 µm² (Table 3).

### Quantification of hRTN neurons

Based on Phox2b+/TH− immunoreactivity, the number of putative RTN neurons ranged from 5 to 65 per hemi-section, with a peak observed between Obex +14.0 mm and +14.3 mm (Fig. 7a). Considering both weak and strong Phox2b-ir, the total number of RTN neurons was 5300 and 4903 (n=2 cases, Table 4), spanning 4 mm in rostrocaudal length (Fig. 7a) and ∼1.5-2 mm in dorsoventral length (Fig. 5)

**Fig. 7.**
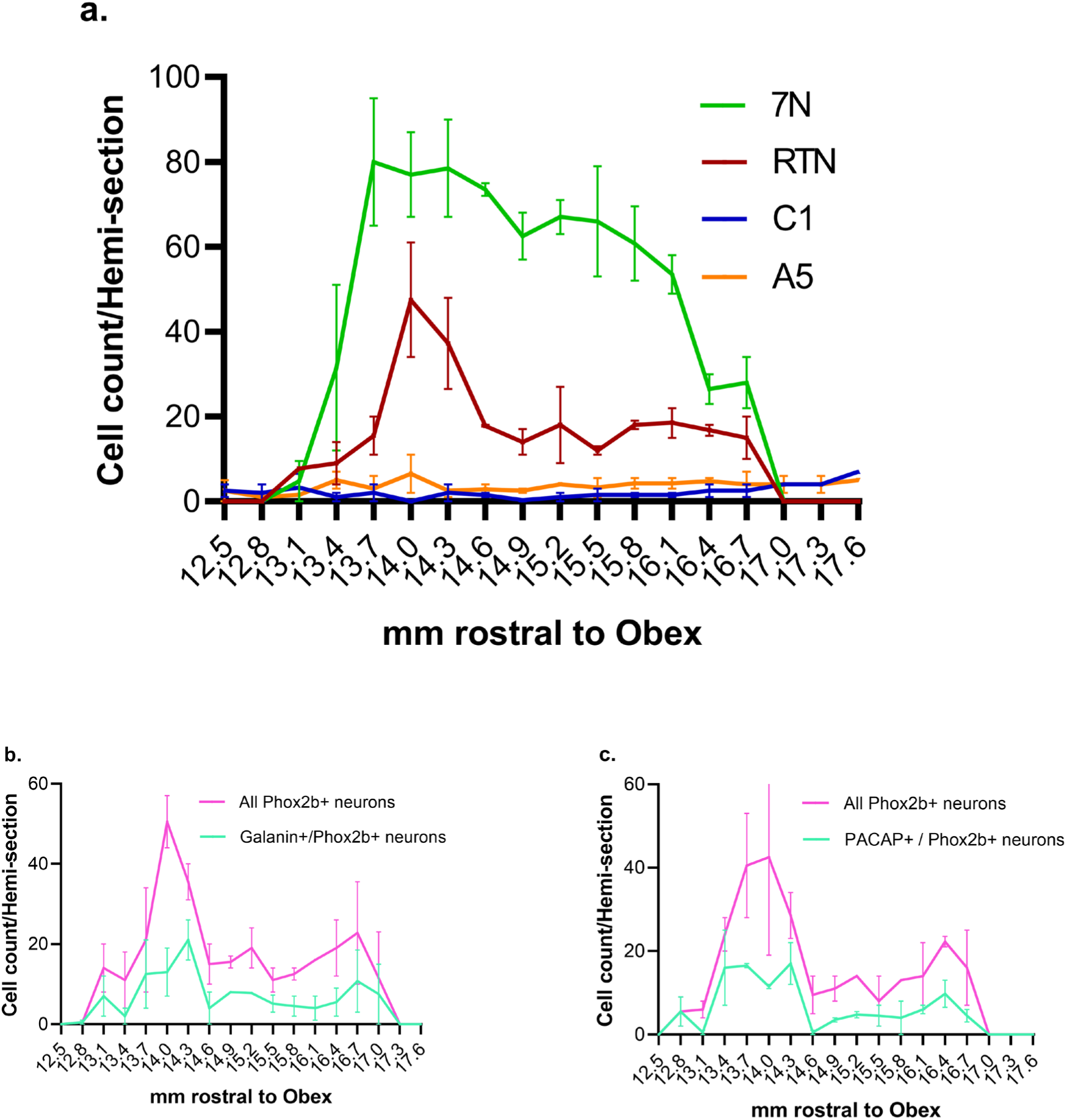
Cell count of the RTN, 7N, C1 and A5 neurons. (a) Rostral-caudal distribution and quantification RTN (Phox2b+/TH−), C1 (Phox2b+/TH+) and A5 (Phox2b−/TH+) neurons within the region of interest specified in Fig. 1b and Fig. 1c. Additionally, total 7N (ChAT+) neurons were counted. The rostral-caudal distribution of (b) Galanin+/Phox2b+ neurons and (c) PACAP+/Phox2b+ neurons, in the RTN region. The curves represent the average number of cells/hemi-section (n=2). The vertical bars indicate interindividual variability in RTN, 7N, C1, and A5 neuron counts observed between the two human cases.

The rostrocaudal distribution of galanin+/Phox2b+ and PACAP+/Phox2b+ parafacial neurons followed the same trend, showing a peak in counts at Obex 14 mm and spanning 4 mm in rostrocaudal length (Fig. 7b and 7c). From these double labelled sections, a total of 5295 Phox2b-ir parafacial neurons were counted (Table 4). The galanin population was 43% of the total Phox2b-ir population while PACAP was 39% (Table 4).

## Discussion

This study provides a comprehensive characterisation of the adult human RTN using histological and neuroanatomical means. Leveraging previous reports from infant, macaque and rodent studies, we identified a large cluster of Phox2b+/TH− neurons ventral to the 7N, aligning with the expected location of the RTN. We present the first in-depth analysis of Phox2b+ RTN neurons, detailing their rostrocaudal distribution, cell counts, morphology and expression of respiratory-related neuropeptides - galanin and PACAP. We also demonstrate puncta from TH+ fibers in close apposition to RTN neurons. These findings establish a crucial baseline for assessing age-related or disease-related changes in the RTN, such as neuronal loss, or altered neuropeptide expression, in respiratory and neurological disorders.

### Adult human RTN neurons are Phox2b positive and subsets express galanin and PACAP immunoreactivity

In this study, the adult hRTN was identified using histological criteria established for identification of RTN neurons in rodents; namely, parafacial neurons that express the transcription factor Phox2b and are devoid of TH and ChAT immunoreactivity (Guyenet and Bayliss, 2022). We identified a highly circumscribed cluster of Phox2b neurons in adult hRTN, in agreement with findings in infants. Approximately 43% of these neurons co-expressed galanin, closely aligning with findings from adult rats (Stornetta et al., 2009). Furthermore, the number of galanin-ir neurons we identified (2,262) is comparable to the ∼2,400 reported in the macaque RTN (Levy et al., 2019). Our study is the first to demonstrate PACAP expression in hRTN, with 40% of Phox2b-ir parafacial neurons co-expressing PACAP. This proportion is substantially lower than what has been reported in adult mice, where PACAP is expressed in 100% of RTN chemoreceptors in neonates and 88% in adulthood (Shi et al., 2017, Shi et al., 2021).

### The location of hRTN neurons in the adult is highly consistent with the distribution reported in human infants and other species

In this study, the spatial distribution of adult hRTN neurons closely aligns with findings from rodents, non-human primates, and human infants (Levy et al., 2019, Levy et al., 2021, Lavezzi et al., 2012, Rudzinski and Kapur, 2010, Shi et al., 2017). Dorsoventrally, the hRTN neurons are situated dorsal to the pontomedullary junction, lateral to the superior olive, and ventrolateral to the 7N. Rostrocaudally, hRTN neurons extended from Obex +13 to +17 mm. A previous study in adults reported PPGAL+/VGlut2+ putative hRTN neurons extending from Obex +14 to +17 mm (Levy et al., 2021). This shorter rostrocaudal extent observed is potentially due to discrepancies between versions of human brainstem atlases. Notably, the Paxinos 2012 atlas used by Levy et al, describes the facial motor nucleus as 3 mm long, while the updated 2020 version used in our study reports it as 4 mm in rostrocaudal length. Nevertheless, both studies demonstrate close proximity between the RTN and 7N throughout their rostrocaudal extent, underscoring the reliability of the findings.

Another key difference between our study and Levy’s (Levy et al., 2021) is the biomarkers used to identify RTN neurons. While our study successfully detected Phox2b-ir, Levy et al. relied on vGlut2 and PPGAL co-expression, however, they did not define the non-galaninergic RTN neurons (Levy et al., 2021) leading to an underestimation of RTN neurons. Thus, our dataset represents the most complete count of RTN neurons to date including both galanin+ and galanin− RTN neurons.

### The adult human contains more RTN neurons than adult rodents and macaque

We identified approximately 5000 Phox2b+/TH− neurons within the putative hRTN, bilaterally, a count significantly higher than the approximately 980 Phox2b-ir neurons reported in human infants (Lavezzi et al., 2012). The discrepancy may stem from differences in immunohistochemistry protocol, or the exclusion of “weakly” Phox2b-ir neurons from the RTN population in earlier studies (Lavezzi et al., 2012, Levy et al., 2019). However, even when counting only strongly Phox2b-ir neurons, we found an average of 2530 per case bilaterally, which is still significantly greater than the 980 reported by Lavezzi et al. (Lavezzi et al., 2012). Since RTN neuron numbers remain stable from late prenatal stages to adulthood in rodents, developmental stage is unlikely the cause. Direct comparisons across studies would require consistent tissue preservation, antigen retrieval and immunohistochemical protocols. Species comparisons report bilateral RTN counts of ∼2,000 in rats (Stornetta et al., 2009, Takakura et al., 2008), ∼700 in mice (Lazarenko et al., 2009, Shi et al., 2017), and ∼2,400 galaninergic neurons in macaques (Levy et al., 2019). Our higher count in humans likely reflects both true interspecies differences and methodological factors such as more extensive serial sectioning and variations in RTN rostrocaudal length across atlases (Paxinos, 2012, Paxinos et al., 2020). Some earlier studies did not assess serial sections and based total cell counts on only 1-2 sections per case (Lavezzi et al., 2012, Rudzinski and Kapur, 2010). Notably, our female case had 29% more RTN neurons than the male, suggesting possible sex or inter-individual differences that warrant further investigation.

### C1-RTN circuit and its role in breathing control: evidence of neuronal interactions

In this study, we show that TH-ir presumptive C1 neurons are intermingled with RTN neurons, and our data suggest close appositions between TH-ir terminals and RTN neurons. Rodent studies have demonstrated that the RTN is dense with PNMT-ir terminals (Rosin et al., 2006) and vigorously activated by adrenaline via α1 adrenoceptors (Oliveira et al., 2016). Furthermore, optogenetic stimulation of C1 neurons in rats produces respiratory effects similar to RTN photo-stimulation (Souza et al., 2020), suggesting a functional C1-RTN circuit in the central control of breathing. These findings suggest that hRTN neurons receive projections from C1 catecholaminergic neurons, however, further studies are needed to confirm the origin of the TH-ir axons and boutons apposing hRTN neurons.

### Factors affecting RTN neuronal counts and future research directions

The RTN neuronal cell count may be influenced by several factors. First, the neurochemical signature used to define RTN neurons could inadvertently include Phox2b+/TH− PCRt, IRt or Sp5O neurons located in the region dorsal and medial to 7N (Fig. 3d), However these were excluded from hRTN because their distribution was anatomically discontinuous from the RTN cluster. Secondly, although this study identified hRTN neurons based on the neurochemical signature established in rodent functional studies (Lazarenko et al., 2009), further research is needed to confirm that these Phox2b+/TH− neurons are indeed chemosensory (Shi et al., 2017). Third, we have reported RTN neuron counts in older adults, and it is uncertain whether Phox2b-ir neurons undergo age-related degeneration. If such degeneration occurs, our quantitative results may underestimate the number of Phox2b-ir RTN neurons in young adults. Finally, although this study was limited by a small sample size, a key strength lies in our thorough analysis of the entire length of the RTN (14 sections per case, spanning 4mm rostrocaudally), rather than focussing on a single plane as is common in studies using post-mortem human brain tissue. A further strength is the consistent rostral-caudal distribution and the quantified peaks of RTN neurons observed across the two cases. Future studies with a larger sample size are needed to evaluate changes in this neuronal population with age, sex and disease, strengthening the robustness and generalisability of our findings. Our study lays the groundwork for these future investigations and for future studies investigating changes in disorders affecting central respiratory chemoreception.

## Conclusion

This study mapped the full extent of the adult hRTN using established neurochemical and anatomical markers, locating it ventral to 7N, lateral to the superior olive, and overlapped with C1 or A5 catecholaminergic neurons. Extending rostrocaudally from Obex +13 to +17 mm, the hRTN contains around 5000 neurons bilaterally - comprising 90% of Phox2b-ir neurons in the parafacial area - surrounded by dense TH-ir fibers. Approximately 40% of these neurons express galanin and PACAP. The data herein provides baseline values for future studies in clinical contexts such as CCHS and sleep apnoea.

## Supporting information

Supplementary section

## Acknowledgements

This work was supported by grants from Australian Research Council (grant no. DP180101890). Tissues were received from the Sydney Brain Bank which is supported by Neuroscience Research Australia and a special gift in memory of Jim Raftos from the Shaw family. We thank Prof Patrice G Guyenet and Prof Ruth Stornetta for providing feedback on an advanced version of the manuscript.

## Abbreviations

4V: 4th ventricle
6N: abducens nucleus
6n: abducens nerve
7N: facial motor nucleus
7n: facial nerve
8n: vestibulocochlear nerve
A5: A5 catecholaminergic neurons
Biotin-SP-conjugated: biotin-succinimidyl ester conjugated
C1: C1 adrenergic neurons
CCHS: congenital hypoventilation syndrome
ChAT: choline acetyltransferase
CO₂: carbon dioxide
D: dorsal
DAB: 3,3’-diaminobenzidine
DPX: dibutylphthalate polystyrene xylene
FFPE: formalin-fixed, paraffin-embedded
H₂O₂: hydrogen peroxide
H&E: hematoxylin & eosin
hRTN: human retrotrapezoid nucleus
HIER: heat-induced epitope retrieval
icp: inferior cerebellar peduncle
IC: inferior colliculus
IF: immunofluorescence
IgG: immunoglobulin G
IHC: immunohistochemistry
IO: inferior olive
IOP: inferior olive, principal nucleus
-ir: immunoreactivity
IRt: intermediate reticular nucleus
IS: inferior salivatory nucleus
ISH: in situ hybridisation
mcp: medial cerebellar peduncle
ml: medial lemniscus
MSO: medial superior olive
NDS: normal donkey serum
NK1R: neurokinin-1 receptor
NMB: neuromedin B
NTS: nucleus of the solitary tract
PACAP: pituitary adenylate cyclase activating polypeptide
PBS: phosphate-buffered saline
PCRt: parvocellular reticular nucleus
PCRtA: parvocellular reticular nucleus, alpha
Phox2b: paired-like homeobox 2B
PPGAL: preprogalanin
Pn: pontine nuclei
py: pyramidal tract
R: rostral
Rn: Red nucleus
RNA: ribonucleic acid
ROI: regions of interest
RRID: Research Resource Identifier
RTN: retrotrapezoid nucleus
SC: superior colliculus
SLC17A6: solute carrier family 17 member A6 (vesicular glutamate transporter 2 gene)
SO: superior olive
Sp5O: spinal trigeminal nucleus, oral part
sp5: spinal trigeminal tract, nerve
TBS: Tris-buffered saline
TH: tyrosine hydroxylase
Thal: thalamus
V: ventral
vGlut2: vesicular glutamate transporter 2
VRC: ventral respiratory column
WB: Western blotting.

## Author Contributions

This study was designed by YL, CES, RM and NNK. IM trained YL in tissue preparation and immunohistochemistry on FFPE tissues. All authors contributed to the methodology validation and experiment troubleshooting. The tissue preparation, experiments, data collection, data analysis and first manuscript draft were produced by YL. NNK, YL, RM and CES revised and modified manuscript drafts.

## Data Availability

The datasets generated during the current study are available from the corresponding author on reasonable request.

## Ethics approval

Donor tissue was previously collected by the Sydney Brain Bank under UNSW ethics project no. HC#200026. This research project (Project no. HC#200901) was approved by the Human Research Ethics Advisory Panel of the University of New South Wales.

## Consent to participate

Informed consent was obtained from individual participants included in the study prior to tissue donation.

## Consent to publish

The authors affirm that human research participants provided informed consent for use of their data for publication.

---

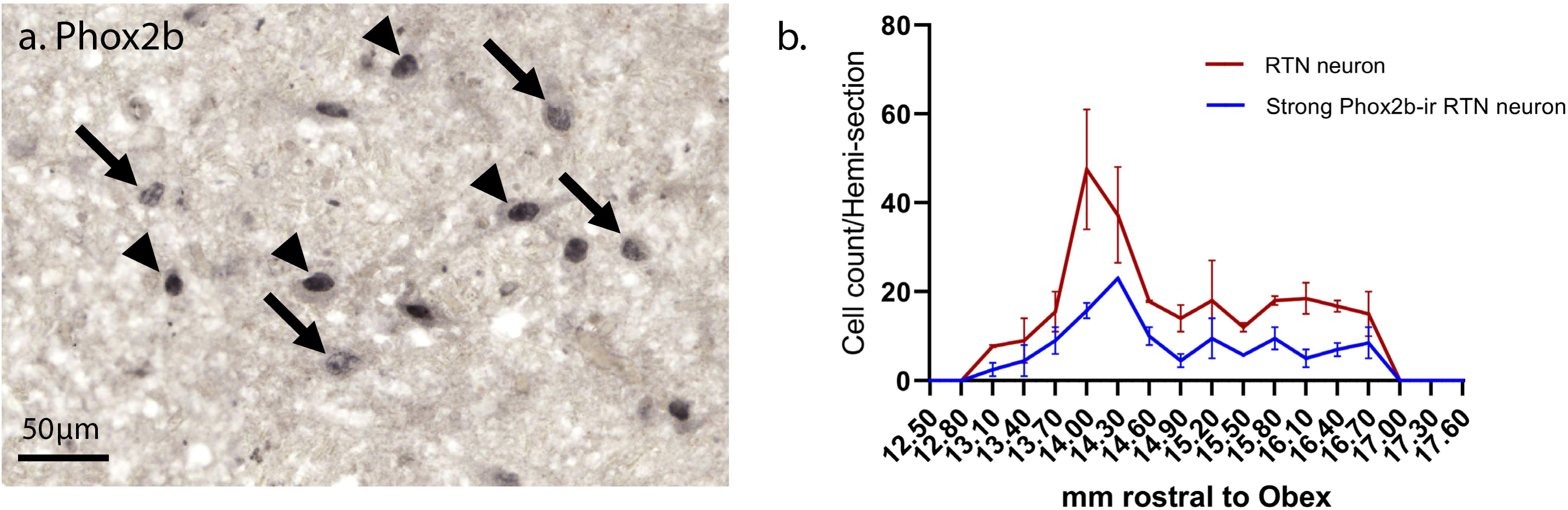

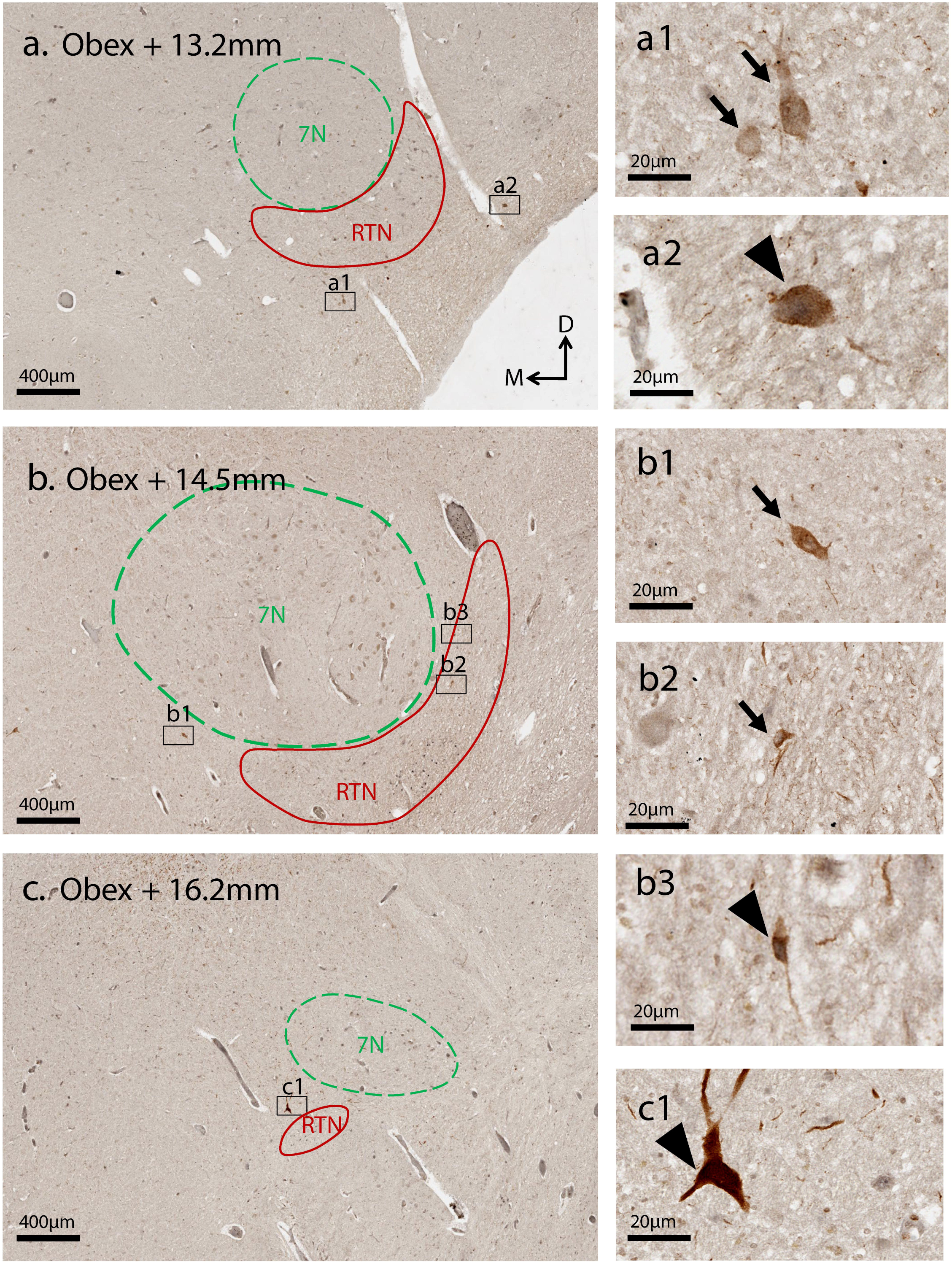

## References

Abbott, S. B., Burke, P. G. & Pilowsky, P. M. 2009. Galanin microinjection into the PreBotzinger or the Botzinger Complex terminates central inspiratory activity and reduces responses to hypoxia and hypercapnia in rat. Respir Physiol Neurobiol, 167, 299–306.10.1016/j.resp.2009.06.003

Abercrombie, M. 1946. Estimation of nuclear population from microtome sections. The Anatomical Record, 94, 239–247.10.1002/ar.1090940210

Agostinelli, L. J., Seaman, S. C., Saper, C. B., et al. 2023. Human Brainstem and Cerebellum Atlas: Chemoarchitecture and Cytoarchitecture Paired to MRI. J Neurosci, 43, 221–239.10.1523/Jneurosci.0587-22.2022

Amiel, J., Dubreuil, V., Ramanantsoa, N., et al. 2009. PHOX2B in respiratory control: lessons from congenital central hypoventilation syndrome and its mouse models. Respir Physiol Neurobiol, 168, 125–32.10.1016/j.resp.2009.03.005

Bankhead, P., Loughrey, M. B., Fernandez, J. A., et al. 2017. QuPath: Open source software for digital pathology image analysis. Sci Rep, 7, 16878.10.1038/s41598-017-17204-5

Bochorishvili, G., Stornetta, R. L., Coates, M. B., et al. 2012. Pre-Bötzinger complex receives glutamatergic innervation from galaninergic and other retrotrapezoid nucleus neurons. J Comp Neurol, 520, 1047–61.10.1002/cne.22769

DEGL’innocenti, D., Becatti, M., Peruzzi, M., et al. 2018. Systemic oxidative stress in congenital central hypoventilation syndrome. Eur Respir J, 52.10.1183/13993003.01497-2018

Dereli, A. S., Oh, A. Y. S., Mcmullan, S., et al. 2024. Galaninergic and hypercapnia-activated neuronal projections to the ventral respiratory column. Brain Struct Funct, 229, 1121–1142.10.1007/s00429-024-02782-8

Dereli, A. S., Yaseen, Z., Carrive, P., et al. 2019. Adaptation of Respiratory-Related Brain Regions to Long-Term Hypercapnia: Focus on Neuropeptides in the RTN. Front Neurosci, 13, 1343.10.3389/fnins.2019.01343

Goridis, C. & Brunet, J. F. 2010. Central chemoreception: lessons from mouse and human genetics. Respir Physiol Neurobiol, 173, 312–21.10.1016/j.resp.2010.03.014

Guyenet, P. G. & Bayliss, D. A. 2015. Neural Control of Breathing and CO2 Homeostasis. Neuron, 87, 946–61.10.1016/j.neuron.2015.08.001

Guyenet, P. G. & Bayliss, D. A. 2022. Central respiratory chemoreception. Handb Clin Neurol, 188, 37–72.10.1016/b978-0-323-91534-2.00007-2

Guyenet, P. G., Stornetta, R. L., Souza, G., et al. 2019. The Retrotrapezoid Nucleus: Central Chemoreceptor and Regulator of Breathing Automaticity. Trends Neurosci, 42, 807–824.10.1016/j.tins.2019.09.002

Huang, J., Waters, K. A. & Machaalani, R. 2017. Pituitary adenylate cyclase activating polypeptide (Pacap) and its receptor 1 (PAC1) in the human infant brain and changes in the Sudden Infant Death Syndrome (SIDS). Neurobiol Dis, 103, 70–77.10.1016/j.nbd.2017.04.002

Ikeda, K., Kawakami, K., Onimaru, H., et al. 2017. The respiratory control mechanisms in the brainstem and spinal cord: integrative views of the neuroanatomy and neurophysiology. J Physiol Sci, 67, 45–62.10.1007/s12576-016-0475-y

Kang, B. J., Chang, D. A., Mackay, D. D., et al. 2007. Central nervous system distribution of the transcription factor Phox2b in the adult rat. J Comp Neurol, 503, 627–41.10.1002/cne.21409

Kasi, A. S., Kun, S. S., Keens, T. G., et al. 2018. Adult With PHOX2B Mutation and Late-Onset Congenital Central Hypoventilation Syndrome. J Clin Sleep Med, 14, 2079–2081.10.5664/jcsm.7542

Kordower, J. H., Le, H. K. & Mufson, E. J. 1992. Galanin immunoreactivity in the primate central nervous system. J Comp Neurol, 319, 479–500.10.1002/cne.903190403

Lavezzi, A. M., Weese-Mayer, D. E., Yu, M. Y., et al. 2012. Developmental alterations of the respiratory human retrotrapezoid nucleus in sudden unexplained fetal and infant death. Auton Neurosci, 170, 12–9.10.1016/j.autneu.2012.06.005

Lazarenko, R. M., Milner, T. A., Depuy, S. D., et al. 2009. Acid sensitivity and ultrastructure of the retrotrapezoid nucleus in Phox2b-EGFP transgenic mice. J Comp Neurol, 517, 69–86.10.1002/cne.22136

Levy, J., Droz-Bartholet, F., Achour, M., et al. 2021. Parafacial neurons in the human brainstem express specific markers for neurons of the retrotrapezoid nucleus. J Comp Neurol, 529, 3313–3320.10.1002/cne.25191

Levy, J., Facchinetti, P., Jan, C., et al. 2019. Tridimensional mapping of Phox2b expressing neurons in the brainstem of adult Macaca fascicularis and identification of the retrotrapezoid nucleus. J Comp Neurol, 527, 2875–2884.10.1002/cne.24713

Meylemans, A., Depuydt, P., De Baere, E., et al. 2021. Adult-onset congenital central hypoventilation syndrome due to PHOX2B mutation. Acta Neurol Belg, 121, 23–35.10.1007/s13760-020-01363-w

Oliveira, L. M., Moreira, T. S., Kuo, F. S., et al. 2016. alpha1- and alpha2-adrenergic receptors in the retrotrapezoid nucleus differentially regulate breathing in anesthetized adult rats. J Neurophysiol, 116, 1036–48.10.1152/jn.00023.2016

Paxinos, G., Furlong, T. & Watson, C. 2020. Human brainstem: cytoarchitecture, chemoarchitecture, myeloarchitecture, Academic Press.

Paxinos, G., Xu-Feng, H., Sengul, G., & Watson, C 2012. Organization of brainstem nuclei. In The human nervous system Elsevier, 260–327

Rosin, D. L., Chang, D. A. & Guyenet, P. G. 2006. Afferent and efferent connections of the rat retrotrapezoid nucleus. J Comp Neurol, 499, 64–89.10.1002/cne.21105

Rudzinski, E. & Kapur, R. P. 2010. PHOX2B immunolocalization of the candidate human retrotrapezoid nucleus. Pediatr Dev Pathol, 13, 291–9.10.2350/09-07-0682-Oa.1

Shi, Y., Stornetta, D. S., Reklow, R. J., et al. 2021. A brainstem peptide system activated at birth protects postnatal breathing. Nature, 589, 426–430.10.1038/s41586-020-2991-4

Shi, Y., Stornetta, R. L., Stornetta, D. S., et al. 2017. Neuromedin B Expression Defines the Mouse Retrotrapezoid Nucleus. J Neurosci, 37, 11744–11757.10.1523/Jneurosci.2055-17.2017

Souza, G., Stornetta, R. L., Stornetta, D. S., et al. 2020. Differential Contribution of the Retrotrapezoid Nucleus and C1 Neurons to Active Expiration and Arousal in Rats. J Neurosci, 40, 8683–8697.10.1523/jneurosci.1006-20.2020

Stornetta, R. L., Spirovski, D., Moreira, T. S., et al. 2009. Galanin is a selective marker of the retrotrapezoid nucleus in rats. J Comp Neurol, 512, 373–83.10.1002/cne.21897

Takakura, A. C., Moreira, T. S., Stornetta, R. L., et al. 2008. Selective lesion of retrotrapezoid Phox2b-expressing neurons raises the apnoeic threshold in rats. J Physiol, 586, 2975–91.10.1113/jphysiol.2008.153163

