## Supplementary section for "Characterisation of putative retrotrapezoid nucleus (RTN) chemoreceptor neurons in the adult human brainstem"

**Table of Contents**

Supplementary Table 1: Demographic and postmortem characteristics of study cases

Supplementary Table 2. Tissue sequence for immunohistochemical labelling of adjacent 10um sections

Supplementary Material 1: Optimisation of Phox2b Immunohistochemistry (IHC) in Human FFPE Tissue

Supplementary Material 2: Using QuPath for cell quantification

Supplementary Material 3: Total RTN cell count and stratification by Phox2b labelling intensity

**Supplementary Table 1.** Demographic and postmortem characteristics of study cases

| Case number | Age | Gender | Postmortem delay (hrs) | Time in formalin fixative (months) |
|---|---|---|---|---|
| 1 | 76 | Female | 29 | 34 |
| 2 | 87 | Male | 59 | 34 |

**Supplementary Table 2.** Tissue sequence for immunohistochemical labelling of adjacent 10µm sections

| | H&E stain | ChAT | Phox2b/TH | Phox2b/PACAP | Phox2b/galanin |
|---|---|---|---|---|---|
| **Adjacent sections** | 1 | 2 | 3 | 4 | 5 |
| **Total sections/case** | 18 | 18 | 18 | 18 | 18 |

### Supplementary Material 1: Optimisation of Phox2b immunohistochemistry in human FFPE tissue

Achieving reliable Phox2b-ir in FFPE human brain tissue has proven challenging in previous studies (Table 1 of main manuscript). This difficulty stems from factors such as prolonged tissue fixation duration, inconsistent conditions during postmortem tissue collection, variations in donors' baseline medical conditions and age-related changes in brain tissue (Zhu et al., 2005). To address these challenges and achieve specific and reproducible Phox2b-ir, we optimised the immunohistochemistry protocol by systematically adjusting conditions at each assay step to identify the parameters necessary for reliable Phox2b detection using a mouse anti-Phox2b antibody (sc-376997-X, Santa Cruz Biotechnology, USA). The specificity of this antibody was previously determined in our laboratory in rodent brain tissue through co-localisation with Phox2b mRNA labelling using RNAscope (Dereli et al., 2021).

*1. Tissue Preparation*

a. For optimisation, brain tissue was obtained from cases fixed in formalin for < 5 years, giving priority to those with the shortest fixation duration and post-mortem interval. The pH of the fixation solution was maintained within the appropriate range of 7 to 7.4 to preserve tissue integrity and antigenicity.

b. After sectioning, tissue mounted onto silanised glass slides was air dried for at least 4 hours then at 55°C overnight to improve section adhesion to the slides, which reduces the risk of detachment during antigen retrieval.

*2. Antigen Retrieval*

After several rounds of optimisation, we identified an effective antigen retrieval approach that unmasked the Phox2b antigen with minimal nonspecific immunoreactivity in the parafacial region of human brain tissue fixed in formalin for up to 5 years. Optimisation involved varying retrieval buffers, heating chamber type, time and temperature as detailed below:

a. Retrieval buffers: Citrate buffer (pH 6) yielded the most effective retrieval results when compared to other buffers such as citrate buffer pH 9, 0.05mM EDTA buffer pH 9, and 0.1% Triton X in PBS pH 8.

b. Heating Chamber type and settings: Antigen retrieval performed with a pressure cooker (6 PSI, 110°C, 15 minutes) produced the best results. Shorter retrieval time (e.g., 5, 8 or 10 minutes) or alternative methods (e.g., heated in a microwave to 100°C for 15 minutes, or using a water bath at 95°C for 20 minutes) led to weak or absent Phox2b-ir . For example, antigen retrieval performed in a pressure cooker (6 PSI, 110°C, 5 mins) using citrate buffer pH 6 successfully revealed Phox2b-ir in 1-year fixed

FFPE tissue but failed to reveal Phox2b-ir in 5-year fixed FFPE tissue. Extending antigen retrieval time to 15 mins enabled detection of Phox2b-ir in 5-year FFPE tissue, however, longer durations (e.g., 20 mins) caused significant tissue damage and increased background noise.

*3. Phox2b antibody concentrations*

Titration experiments demonstrated the optimal dilution for the primary antibody as 1:1000, and for the secondary antibody, it was 1:500. This was in comparison to combinations of primary antibody at 1:500 and 1:1500. with secondary at 1:300 and 1:1000 respectively.

**Supplementary Material 2: Using QuPath for cell quantification**

In this study, RTN, 7N, and C1 neurons and A5 neurons were manually quantified using QuPath software (Bankhead et al., 2017). RTN neurons' morphology parameters were analysed with the QuPath "Positive Cell Detection" function. After importing Phox2b/PACAP and Phox2b/galanin dual-labelled images into QuPath, the following detection parameters were applied: Requested pixel size (0.5 µm) → Background radius (8 µm) → Median filter radius (0 µm) → Sigma (1.5 µm) → Minimum area (35 µm^2) → Maximum area (400 µm^2) → Intensity parameter (Threshold 1) → Max background intensity (2) → Split by shape (Yes) → Exclude DAB (No) → Cell expansion (5 µm) → Include cell nucleus (Yes) → Smooth boundaries (Yes) → Make measurements (Yes) → Score compartment (Nucleus: OAB OD mean) → Threshold 1+ (0.2) → Threshold 2+ (0.4) → Threshold 3+ (0.6) → Single threshold (Yes). QuPath automatically calculated the average nucleus area and cell area, from which the cell diameter was derived.

**Supplementary Material 3: Total RTN cell count and stratification by Phox2b labelling intensity**

Phox2b-ir RTN nuclei were either strongly labelled (appearing dark, intense, and grey / black; 48%), or weakly labelled, exhibiting a lighter grey colour, with less distinct nuclear borders (52%) (Supplementary Fig. 1). Both strongly and weakly Phox2b-ir nuclei were clearly distinguishable from the background (Supplementary Fig. 1a). The strongly Phox2b-ir RTN neurons account for approximately half of the total RTN neuron count (Supplementary Fig. 1b and Supplementary Table 3). We observed consistency in tissue series, with counts and percentages of strong and weak Phox2b-ir being similar between double-labelled PACAP/Phox2b and galanin/Phox2b sections (data not shown).

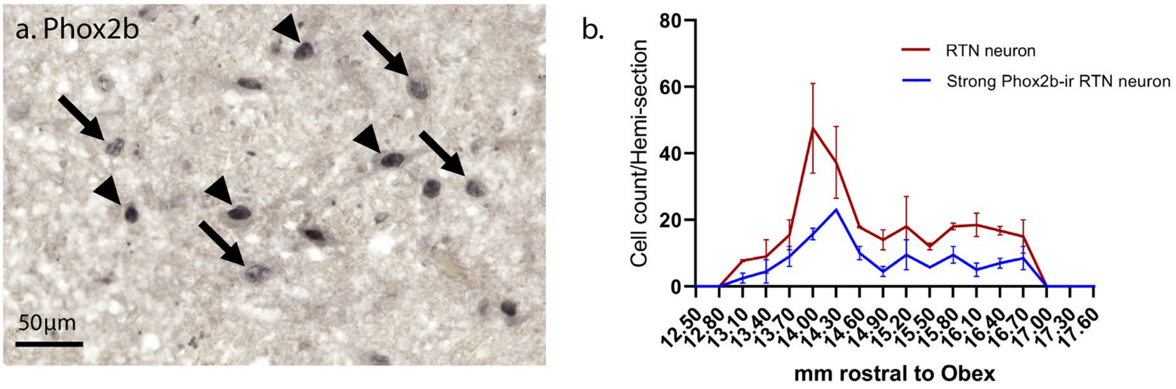


**Supplementary Fig. 1 Stratification of Phox2b labelling intensity in the RTN.** (a) Strong Phox2b-ir neurons are indicated by arrowheads; weak Phox2b-ir neurons are indicated by arrows (b) Rostrocaudal distribution of total RTN neurons and RTN neurons with strong Phox2b-ir labelling intensity in two human cases. The vertical bars indicate interindividual variability in RTN and strong Phox2b-ir RTN counts observed between the two human cases. Due to the small sample size (n = 2), statistical measures such as standard error are not reported

**Supplementary Table 3.** RTN (Phox2b+ / TH-), parafacial (total Phox2b+) neuron counts, stratified by Phox2b labelling intensity and 7N cell count.

| Neuron type | **Phox2b + RTN neurons** | **Strong Phox2b+ RTN (%)** | **Weak Phox2b+ RTN (%)** | **Phox2b + parafacial neurons** | **Strong Phox2b+ parafacial neurons (%)** | **Weak Phox2b+ parafacial neurons (%)** | **7N** |
|---|---|---|---|---|---|---|---|
| **Case 1** | 5300 | 2530 (48%) | 2770 (52%) | 5961 | 2963 (50%) | 2998 (50%) | 9468 |
| **Case 2** | 4903 | 2410 (49%) | 2493 (51%) | 4630 | 2065 (45%) | 2565 (55%) | 6420 |
| **Average** | 5102 | 2470 (48%) | 2632 (52%) | 5296 | 2514 (47%) | 2782 (53%) | 7944 |

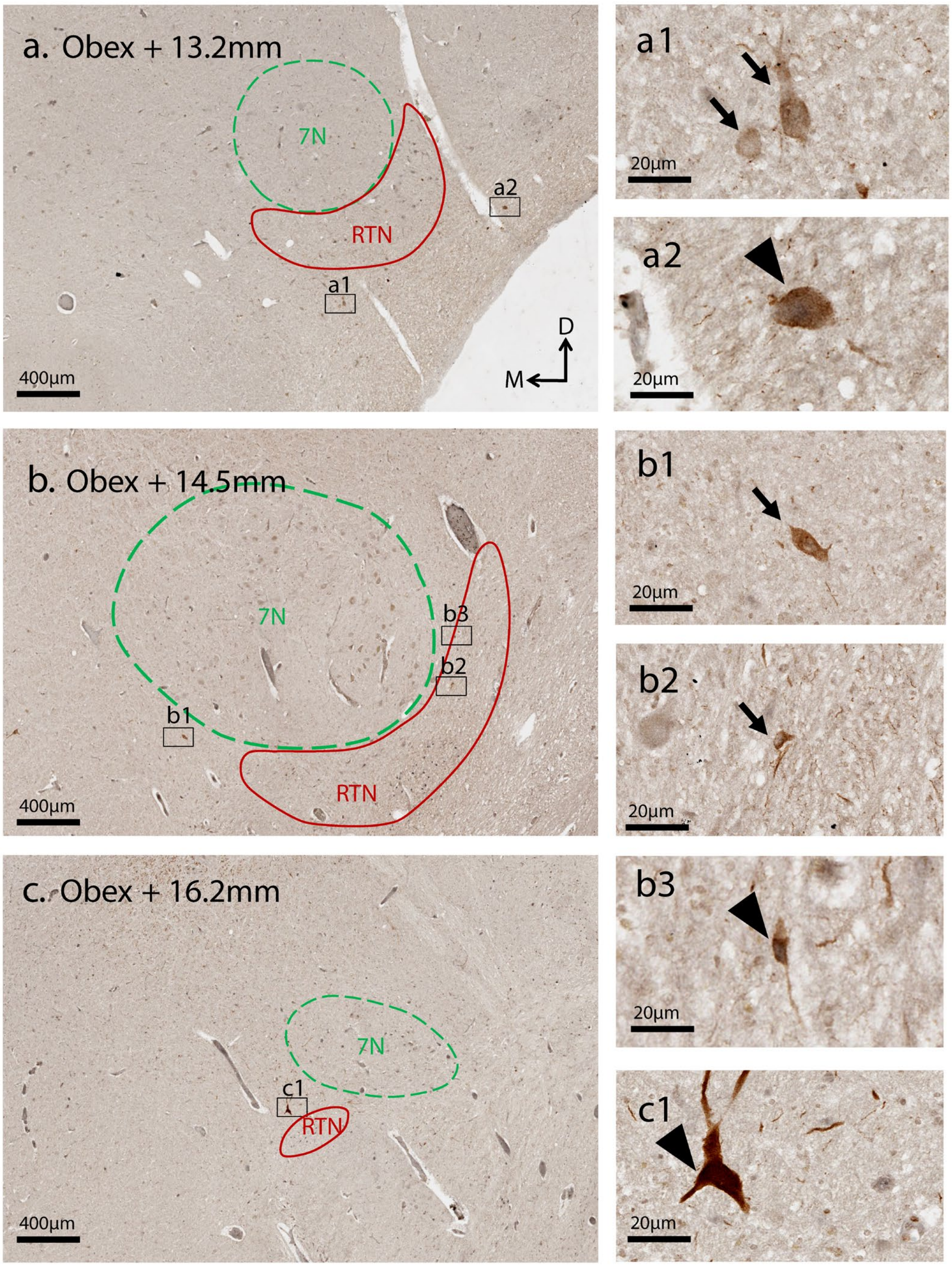


**Supplementary Fig. 2 Distribution of catecholaminergic neurons in the parafacial area at different Obex levels.** (a–c) Low-magnification images showing the parafacial area at different rostrocaudal levels relative to Obex. The red contours indicate the RTN, and the green contours outline the 7N. TH-ir neurons are visible within the parafacial area, representing A5 (Phox2b–/TH+, indicated by arrows) and C1 (Phox2b+/TH+, indicated by arrowheads) neuronal populations. (a1, a2) High-magnification images from Obex +13.2 mm level depicted in (a). In (a1), two A5 neurons are indicated by arrows. In (a2), a single C1 neuron is indicated by an arrowhead. (b1–b3) High-magnification images from regions indicated in (b). In (b1) and (b2), single A5 neurons are marked by arrows. In (b3), a C1 neuron is marked by an arrowhead. (c1) High-magnification image from (c), showing a single C1 neuron indicated by an arrowhead.

**Reference**


BANKHEAD, P., LOUGHREY, M. B., FERNANDEZ, J. A., et al. 2017. QuPath: Open source software for digital pathology image analysis. *Sci Rep,* 7**,** 16878.http://dx.doi.org/10.1038/s41598-017-17204-5

DERELI, A. S., BAILEY, E. J. & KUMAR, N. N. 2021. Combining Multiplex Fluorescence in situ Hybridization with Fluorescent Immunohistochemistry on Fresh Frozen or Fixed Mouse Brain Sections. *J Vis Exp*.http://dx.doi.org/10.3791/61709

ZHU, M. Y., WANG, W. P., IYO, A. H., et al. 2005. Age - associated changes in mRNA levels of Phox2, norepinephrine transporter and dopamine β - hydroxylase in the locus coeruleus and adrenal glands of rats. *Journal of Neurochemistry,* 94**,** 828-838.http://dx.doi.org/10.1111/j.1471-4159.2005.03245.x
